# Attention weights accurately predict language representations in the brain

**DOI:** 10.1101/2022.12.07.519480

**Authors:** Mathis Lamarre, Catherine Chen, Fatma Deniz

## Abstract

In Transformer-based language models (LMs) the attention mechanism converts token embeddings into contextual embeddings that incorporate information from neighboring words. The resulting contextual hidden state embeddings have enabled highly accurate models of brain responses, suggesting that the attention mechanism constructs contextual embeddings that carry information reflected in language-related brain representations. However, it is unclear whether the attention weights that are used to integrate information across words are themselves related to language representations in the brain. To address this question we analyzed functional magnetic resonance imaging (fMRI) recordings of participants reading English language narratives. We provided the narrative text as input to two LMs (BERT and GPT-2) and extracted their corresponding attention weights. We then used encoding models to determine how well attention weights can predict recorded brain responses. We find that attention weights accurately predict brain responses in much of the frontal and temporal cortices. Our results suggest that the attention mechanism itself carries information that is reflected in brain representations. Moreover, these results indicate cortical areas in which context integration may occur.

## 1 Introduction

The attention mechanism has enabled Transformer-based language models (LMs) to achieve state-of-the-art performance on many NLP tasks (Vaswani et al., 2017; Radford et al., 2018; Devlin et al., 2018). In these LMs the attention mechanism iteratively computes weighted sums of the embeddings of neighboring tokens in order to transform token embeddings into contextual representations. The resulting contextual representations have been used to produce predictions of brain responses that are more accurate than predictions derived from (static) lexical embeddings or from RNN-based contextual embeddings (Schrimpf et al., 2021; Toneva and Wehbe, 2019; Caucheteux et al., 2021). These findings show that the attention mechanism produces contextual hidden state embeddings that contain more information reflected in brain responses than lexical embeddings alone. However, prior work has not determined whether the mechanism by which hidden state embeddings are constructed (i.e., the weights computed by the attention mechanism) is itself reflected in brain representations.

To address this question, we used voxelwise encoding models (VM), a powerful data-driven method for determining what information is represented in the brain (Wu et al., 2006; Naselaris et al., 2011). In encoding models, brain responses to stimuli are modeled as a linear combination of feature spaces that each represent properties of the stimuli. These encoding models are used to determine where each feature space is represented in the brain. Encoding models have previously been used to determine how variables such as lexical and contextual semantics are represented in brain responses (Huth et al., 2016; de Heer et al., 2017; Deniz et al., 2019; Jain and Huth, 2018). However, previous work did not directly use attention weights in encoding models to study whether and how the attention weights are reflected in brain responses. Understanding the relationship between attention weights and brain representations would provide insight into where information integration across context occurs in the brain, and whether attention weights contain information relevant for human language processing.

In this work, we use attention weights in encoding models to determine whether weights computed by the attention mechanism of Transformer-based LMs are represented in brain responses. We modeled functional magnetic resonance (fMRI) data recorded from six human participants while they read a set of English natural language narratives. We presented the same stimuli as input to BERT and GPT-2 and extracted their corresponding attention weights. We investigated how well attention weights predict brain responses, compared to other stimulus features such as lexical and contextual semantic word embeddings. We found that, across many cortical areas that have previously been associated with language processing, attention weights better predict brain responses than lexical embeddings. In a subset of these cortical areas, attention weights better predict brain responses than contextual embeddings, suggesting that the mechanism of context integration (attention weights) in Transformer-based LMs reflects aspects of brain responses that are not captured by the resulting contextual embeddings. Moreover, these findings suggest cortical areas that could be involved in the integration of information from context words.

## 2 Related work

### 2.1 NLP features for neurolinguistics

Features extracted from NLP models have been previously used in encoding models to study how language is represented in the human brain. Prior work used lexical word embeddings to show that semantic information is represented across the cerebral cortex (Huth et al., 2016) and that these representations are similar during listening and reading (Deniz et al., 2019). Others used RNNs (Wehbe et al., 2014) or contextual word embeddings (Jain and Huth, 2018) to study where linguistic context is integrated in the brain. Most recently, studies showed that the hidden states of Transformer-based LMs predict brain activity more accurately than lexical embeddings or RNN-based contextual embeddings (Schrimpf et al., 2021; Caucheteux et al., 2021; Toneva and Wehbe, 2019). However, it remains unclear whether the Transformer model’s attention weights contain relevant information to represent language-related information in the brain that is not captured in the resulting contextual embeddings. Recent concurrent work investigated the similarity between brain representations and the activity of the attention heads in BERT (Kumar et al., 2022). From each attention head, the authors extracted the “transformation” vector which is used to update the hidden states. This transformation vector is the sum of the value vectors of each token weighted by the attention weights. They used the transformation vector and its norm as features of encoding models to predict brain responses. Both of these features contain information about the attention weights, but are also dependent on the value vectors, which additionally encode semantic information. In our work, we predict brain responses directly from the attention weights in order to more clearly distinguish between the representations of attention weights and semantic embeddings. In addition, we analyze the attention weights of two Transformer-based LMs (BERT and GPT-2) to determine whether attention weights from bidirectional and unidirectional LMs are reflected differently in brain responses.

### 2.2 Attention analysis in NLP

Several studies have investigated whether the attention mechanism can be analyzed to improve interpretability of Transformer-based LMs (Rogers et al., 2020; Vashishth et al., 2019; Galassi et al., 2020). Because attention defines how much a word will be weighted to compute the next representation of its neighbors, some have claimed that attention is naturally interpretable (Clark et al., 2019). However, whether attention weights actually explain the behavior of LMs has been subject to debate (Jain and Wallace, 2019; Wiegreffe and Pinter, 2019). Nevertheless, there is evidence that attention weights do carry relevant information. At a qualitative level, the network structure of LMs can be used with attention weights to improve interpretability (Abnar and Zuidema, 2020). Moreover, a quantitative analysis of BERT’s 144 attention heads showed that attention weights in some attention heads correspond to linguistic functions such as syntactic or coreference relations (Clark et al., 2019). For example in a particular head, direct objects consistently give the highest attention weight to their verbs. This analysis showed that linguistically specialized heads are primarily found in the middle layers of BERT. Similar findings were reported for GPT-2 (Vig and Belinkov, 2019).

## 3 Methods

To investigate whether attention weights predict brain responses, we analyzed functional magnetic resonance imaging (fMRI) recordings of brain responses to English language narratives (Section 3.1). We used the voxelwise encoding modeling (VM) framework (Wu et al., 2006; Naselaris et al., 2011) to determine where attention weights are reflected in the recorded brain responses (Section 3.2). We identified how well attention weights can predict brain responses relative to lexical semantic features, and compared the performance of attention weights across layers and across two LMs (BERT and GPT-2). Then, to study whether attention weights contain additional information to LM hidden states, we determined how well attention weights extracted from BERT can predict brain responses relative to contextual embeddings extracted from the hidden states of BERT. We further did a focused analysis on cortical regions that have previously been associated with language representations: the high-level auditory cortex (AC), the superior temporal sulcus (STS), Broca’s area and the superior ventral premotor speech area (sPMv) (Fedorenko et al., 2011; Huth et al., 2016; de Heer et al., 2017; Deniz et al., 2019). While the attention mechanism in Transformer-based LMs and the concept of attention in cognitive neuroscience are distinct, the two share a conceptual relationship (Lindsay, 2020). Therefore, we also inspected brain representations in areas of the human attention network: the middle temporal visual area (MT), the intraparietal sulcus (IPS) and the frontal eye field (FEF) (Chen et al., 2019). We visualize the results on cortical maps made with pycortex (Gao et al., 2015).

### 3.1 Recording brain responses to naturalistic language stimuli

Functional magnetic resonance imaging (fMRI) was used to record brain activity in healthy participants while they read a set of English language narratives. fMRI measures the blood-oxygen-level dependent (BOLD) signal, which indirectly measures the intensity of neural activity. Each fMRI image consists of approximately 80,000 cortical voxels, and each voxel corresponds to an approximately 8 mm^3^ cube of the brain. The repetition time (TR) of acquisition is about 2 seconds.

The stimulus narratives were originally recorded for the Moth Radio Hour. The words were presented one-by-one at the center of a screen for the same duration that they are heard in the audio version. Each stimulus narrative contained between 1547 and 3313 words over 10-15 minutes. Overall, the stories contained 24744 total words (2977 unique words). We refer the reader to the original study for details about fMRI data acquisition and preprocessing (Deniz et al., 2019)^1^. In this study we used ten of the stimulus narratives to estimate encoding models, and we used the eleventh held-out narrative to evaluate the estimated models.

### 3.2 Voxelwise encoding models fitting and evaluation

In the VM framework, stimuli are first nonlinearly transformed into sets of features (also called “feature spaces”) hypothesized to be represented in brain activity. These feature spaces *X_i_* ∈ ℝ*^d_i_×t^* are linearly combined to describe brain responses *X_i_* ∈ ℝ*^v×t^*, where *d_i_* is the dimension of each feature space, *t* is the number of timesteps and *v* is the number of voxels. For each feature space *X_i_*, a set of weights *B_i_* ∈ ℝ*^v×d_i_^* is estimated to map from the feature space to brain responses. This set of estimated model weights describes how information in a given feature space modulates BOLD responses in each voxel. In order to account for variance in brain responses that can be explained by multiple feature spaces, the weights for multiple feature spaces are jointly estimated. To jointly estimate these weights, banded ridge regression is used. Banded ridge regression is a variant of standard ridge regression that allows different regularization parameters for each feature space, and therefore can avoid biases caused by differences in feature distributions (Nunez-Elizalde et al., 2019). Mathematically, banded ridge regression estimates the weights for each feature space by solving:

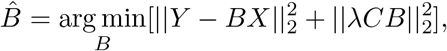

where *C* is a diagonal matrix containing the regularization level. The estimated weights form a *joint encoding model* that describes brain responses as the linear combination of multiple feature spaces. To evaluate the encoding model, the estimated model weights are used to predict brain responses to a held-out stimulus. The prediction performance of the encoding model is computed as the Pearson correlation coefficient (r) between the predicted and recorded brain responses to the held-out stimulus for each voxel. Model prediction performance describes how well the feature spaces can jointly predict the time-course of brain responses. In order to determine how well each individual feature space can predict brain responses, the split-prediction performance is used. The split-prediction performance is a model interpretation metric that decomposes the prediction performance of the joint encoding model into the contribution of each feature space while taking into account variation in brain responses that can be explained by other feature spaces. Formally, for a given feature space, the split-prediction performance *r_i_* is defined as:

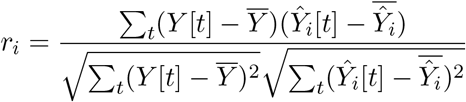

where *Y* is the brain response and *Ŷ_i_* is the prediction of the feature space. This metric has the advantage that it determines the contribution of each feature space in light of other potential explanatory variables, and is more computationally efficient and less conservative than other methods such as variance partitioning. For more details, see St-Yves and Naselaris (2018); la Tour et al. (2022). The statistical significance of the prediction performance r for each voxel is determined by a permutation test with 10000 iterations (Deniz et al., 2019; Lescroart and Gallant, 2019; Jain et al., 2020; Ramakrishnan and Deniz, 2021). In each permutation iteration, the timecourse of voxel responses in the held-out dataset is shuffled in blocks of 10 TRs, and the Pearson correlation is computed between the timecourse of predicted voxel responses and the timecourse of shuffled voxel responses. The resulting correlation coefficients form an empirical null distribution of r for each voxel, which is used to determine the p-value of the observed r for that voxel. This p-value is computed for each voxel separately. A false discovery rate (FDR) correction (Benjamini and Hochberg, 1995) is then applied to the resulting p-values, and voxels with a corrected p-value of < 0.05 are considered to be statistically significant.

### 3.3 Feature spaces

We categorized our variables of interest into three types of feature spaces: attention weights (extracted from BERT or GPT-2), semantic representations (lexical or contextual embeddings), and low-level sensory features. Each type of feature space is detailed in the following subsections.

#### Attention features

The attention feature space reflects the weights computed by the attention mechanism in order to form contextual hidden state representations. We computed the attention feature space either from BERT-base or GPT-2 small (Devlin et al., 2018; Radford et al., 2019). For both models, we used pretrained models available from the HuggingFace library (Wolf et al., 2020). To construct an attention feature space, we first extracted the *attention maps* from the attention heads of the LM. An attention map is a matrix *M* ∈ *R^n×n^,* where *n* is the number of tokens in the input sequence. Each entry *m_i,j_* represents the attention weight from token *i* to token *j*. For each input sequence, we used a sliding window with 10 prior context words, because it has been previously shown to yield to better prediction performance than other context lengths (Toneva and Wehbe, 2019). BERT requires special tokens [CLS] and [SEP] to be appended to the beginning and end of each input sequence. This input procedure resulted in a 11 × 11 matrix M for GPT-2, and a 13 × 13 matrix M for BERT. To convert the attention maps into a feature vector that reflects the average attention to each word in the input sequence, we took the mean over columns of *M*. When extracting attention weights from BERT, we excluded the first and last column that represent the special tokens. For words that are split into multiple tokens during tokenization, we summed over the attention to each of the tokens of the word, following Clark et al., 2019. For each stimulus word, this procedure resulted in a vector of dimension 11 for each attention head. For each layer, the attention feature space contains weights from each of 12 attention heads, resulting in a 132-dimensional feature space. Across all 12 layers, the attention feature space consists of a 1584-dimensional vector for each word.

#### Semantic features

The semantic feature spaces were constructed either from lexical or contextual word embeddings. To construct the lexical semantic feature space we used embeddings from English1000, a 985-dimensional lexical embedding space that has previously been shown to explain variance in brain responses across the cerebral cortex (Huth et al., 2016; Deniz et al., 2019; Vaidya et al., 2022; LeBel et al., 2021; Tang et al., 2021). English1000 is constructed from the co-occurrence statistics in a large corpus of text composed of the Moth stories, 604 popular books available through Project Gutenberg, 2,405,569 Wikipedia pages and 36,333,459 reddit.com user comments. The vocabulary of English1000 includes all the words of the stimulus narratives. To construct the contextual semantic feature space, we used the hidden state activation of the eighth layer of BERT. To construct a contextual embedding of each word, we computed the mean across the embeddings of each token of the word. We used embeddings from the eighth layer of BERT because it produced more accurate predictions of BOLD responses than other layers (Figure 6 in Appendix). This is in accordance with prior work (Schrimpf et al., 2021; Caucheteux et al., 2021; Toneva and Wehbe, 2019).

#### Sensory-level features

To account for brain representations of low-level sensory information, we created eight feature spaces that reflect visual characteristics of the text stimulus: word presentation rate (1 dimension), word length (1 dimension), letter identity (26 dimensions), phoneme rate (1 dimension), phoneme identity (39 dimensions), pauses (1 dimension), word length standard deviation (1 dimension) and motion energy (6555 dimensions). These feature spaces were constructed based on prior work (Huth et al., 2016; de Heer et al., 2017; Deniz et al., 2019). We refer the reader to those studies for additional details.

#### Feature space preprocessing

In order to match the sampling rate of the fMRI recordings, we applied a low-pass Lanczos filter with three lobes with a cut-off frequency at 0.25 Hz to each of the feature spaces (following e.g., Huth et al., 2016; Deniz et al., 2019; Jain and Huth, 2018; Jain et al., 2020). In order to account for the hemodynamic response function of each voxel, which delays the BOLD signal measured by fMRI relative to neural activity, we use a finite impulse response (FIR) model (following Huth et al., 2016; Deniz et al., 2019; Jain and Huth, 2018; Jain et al., 2020). The FIR model was implemented by copying and delaying each feature space by 2, 4, 6, and 8 seconds.

## 4 Results

To study the role of the attention mechanism in explaining brain responses, we tested how well attention weights can predict brain responses to natural language stimuli. We analyzed fMRI recordings that were taken while human participants read a set of English narratives. We provided the text of the stimulus narratives to a Transformer-based language model (BERT or GPT-2) and extracted attention weights from each attention head. We used voxelwise encoding models to determine how well these attention weights can predict brain responses compared to semantic features and low-level sensory features.

### 4.1 Across many brain areas, attention weights explain more variance in brain responses than lexical semantic embeddings

In order to determine how well attention weights extracted from BERT explain brain responses compared to lexical semantic embeddings and sensory-level features, we estimated an encoding model that maps lexical embeddings, BERT attention weights and low-level features to brain responses. In Figure 2, we show the prediction performance of this joint model for one representative participant. Results for five additional participants are qualitatively similar (see Section C in the Appendix). Voxel hue denotes the feature space that best predicted brain response in that voxel: green for attention weights, red for lexical semantic embeddings and blue for low-level features. Voxel opacity denotes the split-prediction performance of the corresponding feature space. The split-prediction performance denotes the contribution of a specific feature space in explaining the variance in the brain response. Voxels shown in gray were not significantly predicted (FDR corrected, *p* < 0.05). Attention weights better predict brain responses in voxels located across the temporal and prefrontal cortices that have previously been associated with language processing: high-level AC, STS, Broca’s area and sPMv. These results suggest that, in many language-related cortical areas, brain representations reflect more information about integration across words, than the lexical representation of the words themselves.

**Figure 1:**
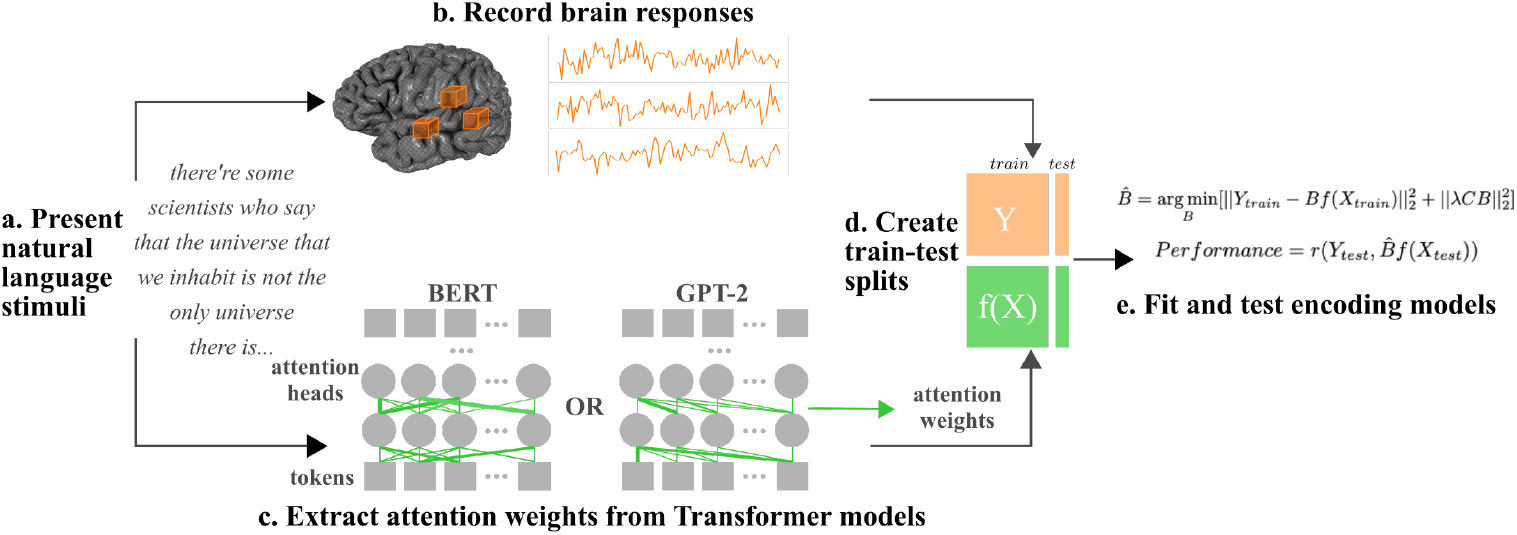
Voxelwise encoding modeling approach. (a) Natural language narratives were used as stimuli. b) Brain responses were recorded using functional MRI while participants read the stimuli. (c) Attention feature spaces were constructed by extracting attention weights from Transformer-based language models, BERT or GPT-2. Additional feature spaces (not shown) were constructed to reflect semantic information and low-level sensory features. (d) Separate train and test sets were created from the feature spaces and from the brain responses. (e) Regularized regression was used to fit an encoding model that predicts brain responses from the extracted feature spaces, in each voxel and participant separately. Model prediction performance was quantified as the Pearson correlation coefficient between predicted brain responses and the recorded brain responses of the test set.

**Figure 2:**
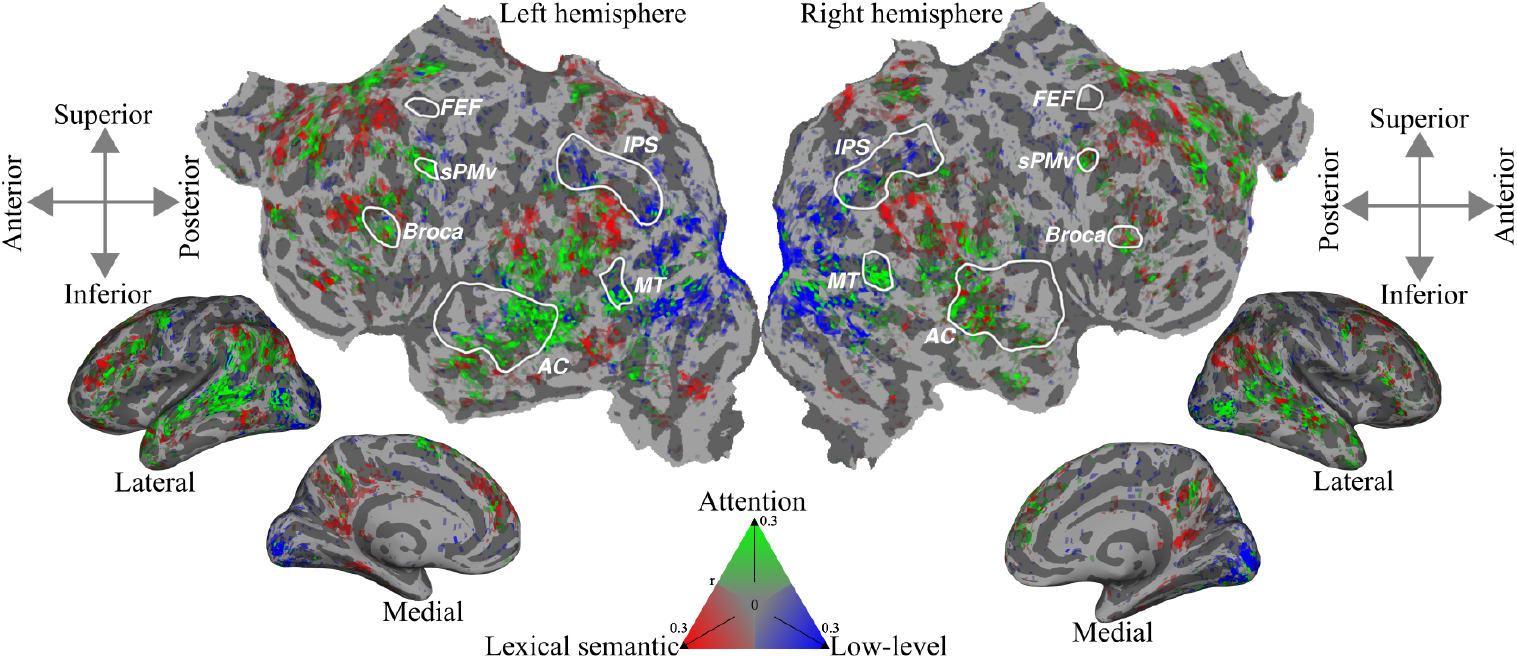
Prediction performance of BERT attention weights, lexical semantic and low-level features across the cerebral cortex. Attention weights extracted from BERT, lexical embeddings, and low-level sensory features were jointly used in an encoding model to predict brain responses. Voxel hue indicates the feature space that best explains voxel responses (green for attention weights, red for semantic features, blue for low-level features). Voxel opacity indicates the split-prediction performance of the best performing model. Voxels shown in gray were not significantly predicted (p < 0.05, FDR-corrected). Attention weights best predict brain responses in most frontal and temporal language-related regions, including Broca’s area, high-level auditory cortex (AC), the superior temporal sulcus (STS), superior ventral premotor speech area (sPMv), and the middle temporal visual area (MT). Lexical embeddings predict best in some frontal and parietal areas. Low-level features predict well mainly in the early visual cortex and the intraparietal sulcus (IPS).

### 4.2 Attention weights from middle layers of BERT explain the most variance in brain responses

In order to determine how well attention weights from different layers of BERT represent information in brain responses, we computed the prediction performance of attention weights extracted from each layer. For each layer, we estimated an encoding model mapping from attention weights in that layer, lexical embeddings and low-level features to brain responses. In Figure 3, we show the prediction performance of attention weights in this layer-wise comparison. The dotted green line shows the prediction of each layer averaged across participants. Each cross (“x”) represents one participant. Attention weights extracted from intermediate layers (layers 5 to 10) best predict brain responses. Subsequently, we compared encoding models with two equally-sized groups of layers: an encoding model with the attention weights of the six intermediate layers grouped (layers 5 to 10, intermediate layers model) and an encoding model with the six early (1 to 4) and late layers (11 to 12) grouped (early and late layers model). The intermediate layers model (dash-dotted green line) performs almost as well as the full model with all the layers (solid green line). The early and late layers model (dashed green line) performs worse than the full model with all the layers (solid green line). The fact that using only half of the layers hardly hurts the performance of the model is consistent with literature on BERT pruning (Voita et al., 2019; Michel et al., 2019). Moreover, the intermediate layers are the ones where linguistically relevant attention heads were found (Clark et al., 2019). In contrast, the attention heads in early and late layers were shown to mostly attend uniformly to every token in the input sequence. This suggests that the attention weights that are linguistically relevant are better at predicting brain responses.

**Figure 3:**
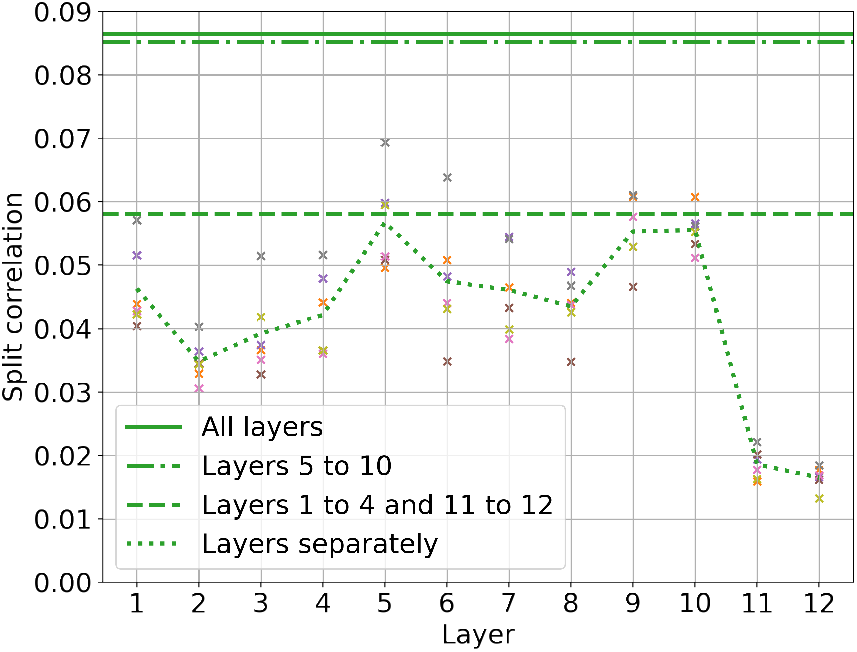
Layer-wise comparison of BERT attention weights prediction performance. To determine how well attention weights in different layers of BERT predict brain responses, a separate encoding model was estimated for each layer. Each encoding model consisted of the attention weights from one layer of BERT, lexical embeddings, and low-level features. The split-prediction performance of the attention features is shown for each encoding model. The dotted line shows the performance of attention weights from each layer separately. Each cross (“x”) shows the per-layer performance for one participant. The solid line shows the prediction performance of attention weights from all 12 layers together. The dash-dotted line shows the performance of attention weights from middle layers (5 to 10) only. The dashed line shows the performance of attention weights from early (1 to 4) and late (11 to 12) layers only. Attention weights extracted from layers 5 to 10 best predict brain responses, suggesting that the attention weights in middle layers carry more brain-relevant information than attention weights in other layers.

### 4.3 The attention weights of BERT and GPT-2 accurately predict brain responses

In order to determine whether the contribution of attention in explaining brain responses is consistent across Transformer-based LMs, we additionally estimated an encoding model using features extracted from GPT-2 attention weights, lexical embeddings and low-level features. In Figure 4, we show the split-prediction performance of GPT-2 attention weights, lexical embeddings and sensory-level features for selected cortical regions. We also show the split-prediction performance of a joint model with BERT attention weights, lexical embeddings and sensory-level features. Each cross (“x”) represents a participant and the height of the bar is the average across participants. The attention weights of both BERT and GPT-2 accurately predict brain responses in the language-related areas such as high-level AC, Broca’s area and sPMv. Interestingly, attention weights predict brain responses also well in the visual attention area MT. However, brain responses in other areas of the attention circuit such as FEF or IPS are not well predicted. Overall, the prediction performance of BERT and GPT-2 attention weights are very similar. This similarity suggests that when only the preceding context is provided to a LM, attention weights from unidirectional and bidirectional models may perform similarly well in terms of cognitive fit.

**Figure 4:**
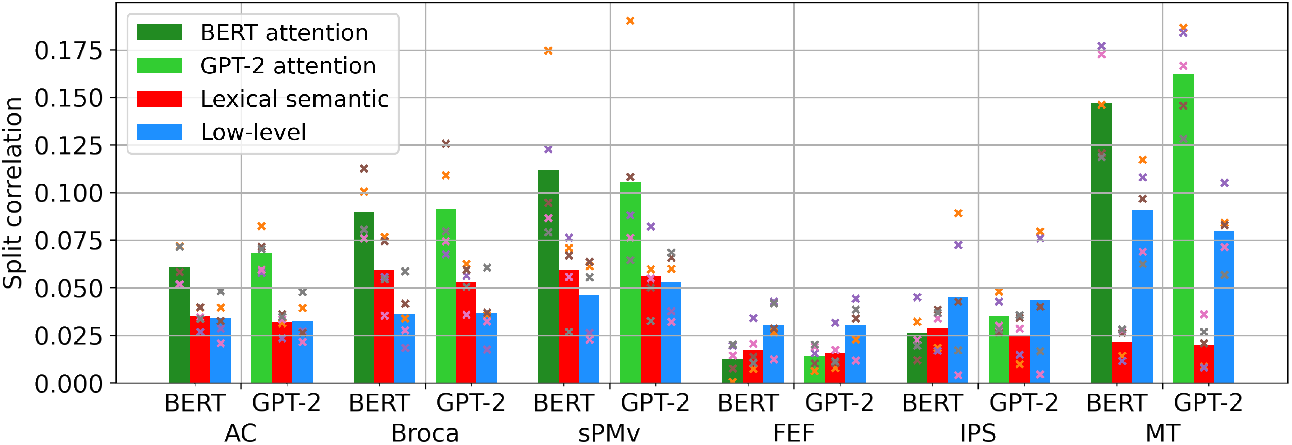
Region-level comparison of BERT and GPT-2 attention weights. We fit two encoding models, each consisting of attention weights, lexical embeddings, and low-level features. We used attention weights from BERT in one encoding model, and from GPT-2 in the other encoding model. The split-prediction performance of each feature space is shown for each LM and cortical region. Average split-prediction performance of attention weights across participants is shown by green bars (dark green for BERT and light green for GPT-2). Average split-prediction performance of lexical embeddings across participants is shown by red bars. Average split-prediction performance of low-level features across participants is shown by blue bars. Each cross “x” represents one participant. BERT and GPT-2 perform very similarly in all regions. Attention weights predict well in language-related regions (AC, Broca’s, sPMv) and in the visual attention area MT.

### 4.4 Across specific temporal and frontal areas, attention weights explain more variance in brain responses than contextual embeddings

Contextual embeddings in the hidden states of Transformer-based LMs carry information about attention weights as well as lexical embeddings. Prior work showed that these hidden state embeddings produce highly accurate predictions of brain responses (Schrimpf et al., 2021; Toneva and Wehbe, 2019). To determine whether attention weights capture information reflected in brain representations that is not present in the contextual embeddings, we compared the prediction performance of attention weights and of hidden state embeddings. We used hidden state embeddings from the eighth layer of BERT, because hidden state embeddings from this layer produced more accurate predictions of brain responses than other layers (see Figure 6 in Appendix). We estimated an encoding model mapping from contextual embeddings from the eighth layer of BERT, BERT attention weights and low-level features to brain responses. In Figure 5, we show the split-prediction performance of each feature space for one representative participant. Results for five additional participants were qualitatively similar and are shown in the Section C of the Appendix. Voxel hue denotes the feature space that best predicted brain response in that voxel: green for attention weights, red for contextual embeddings and blue for low-level features. Voxel opacity denotes the split-prediction performance of the corresponding feature space. Voxels shown in gray were not significantly predicted. Contextual embeddings best predict brain responses in the majority of temporal and prefrontal areas. Yet, attention weights explain most of the brain responses in some of these areas, such as portions of high-level AC, Broca’s area and sPMv. This supports findings that linguistic information in BERT can be found not only in the hidden states, but also in the attention weights (Clark et al., 2019).

**Figure 5:**
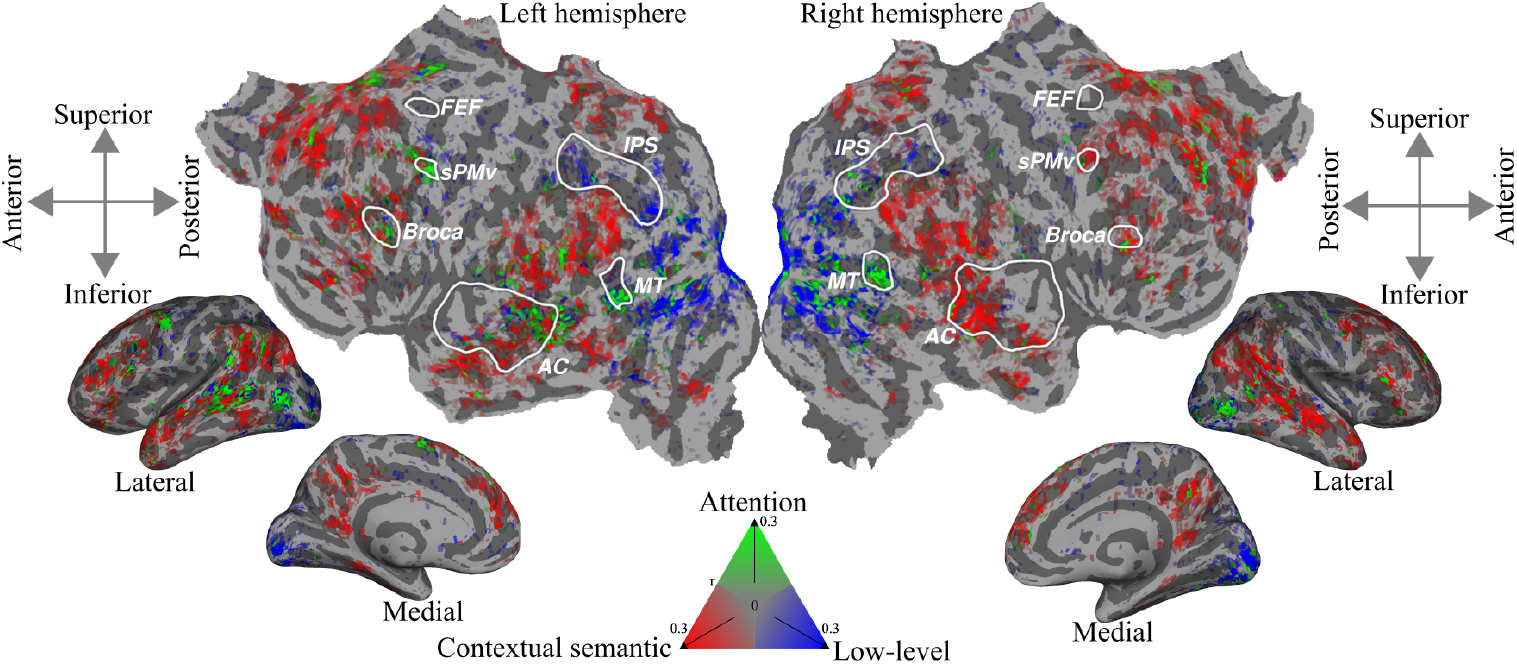
Prediction performance of BERT attention weights, contextual embeddings and low-level features across the cortex. BERT attention weights, contextual semantic embeddings (hidden state embeddings from BERT layer 8), and low-level features were used to estimate an encoding model. Voxel hue indicates the feature space that best explains voxel responses (green for attention weights, red for semantic features, blue for low-level features). Voxel opacity indicates the split-prediction performance of the best performing feature space. Voxels shown in gray were not significantly predicted (*p* < 0.05, FDR-corrected). The attention features best predict some voxels in high-level AC along superior temporal sulcus, Broca’s area, and sPMv. Most MT voxels are well predicted by the attention weights. Contextual embeddings best predict voxels in most frontal, temporal and parietal areas. Low-level features predict best in visual cortex. In most cortical areas, contextual embeddings better predict brain responses than attention weights, but attention weights predict better in some language-related regions and in visual area MT.

## 5 Conclusion

In this work, we examined how well attention weights from Transformer-based LMs predict brain responses to linguistic stimuli. We find that attention weights from BERT and GPT-2 produce accurate predictions across many cortical areas that have previously been associated with language processing (high-level auditory cortex, superior temporal sulcus, Broca’s area, superior ventral premotor speech area). Attention weights better predict brain responses than lexical embeddings in most of these areas. Attention weights are also better at predicting brain responses than contextual embeddings in portions of the language related areas. These findings show that attention weights capture information that is not contained in lexical or contextual embeddings, and indicate cortical areas that could be involved in integrating context across words. Moreover, attention weights from the middle layers of the LM, that have been previously identified as linguistically relevant, contribute strongest to the predictions. While our results showed consistency between unidirectional and bidirectional models in terms of how well attention weights predict brain responses, more work is needed to understand the impact of model directionality on cognitive fit. In the future, comparing the results of our method of computing attention weights with more cognitively-inspired methods could provide more specific insights into how attention weights in Transformer-based LMs relate to processes in the human brain.

### Limitations

Some limitations of our work arise from the nature of fMRI data. The temporal resolution of fMRI (2s) does not capture all the rapid changes in human speech. Using encoding models with another neuroimaging technique with higher temporal resolution, such as MEG, could provide complementary results. Our analyses were limited to English language comprehension, but it could be the case that the nature of contextual language integration differs between different languages. In the future, we hope to apply our methods to other languages in order to obtain a more comprehensive understanding of context integration in the brain and in artificial neural networks. Lastly, note that our analyses demonstrate the similarity of representations between Transformer-based LMs and brain responses, but we caution that these analyses do not directly prove the existence of similar underlying computational mechanisms.

## Acknowledgements

We thank Leila Wehbe for valuable discussions. This work was funded by grants from the National Science Foundation (NAT-1912373) and the German Federal Ministry of Education and Research (BMBF 01GQ1906). CC was supported in part by an NSF GRFP and an IBM PhD fellowship.

## A Computing infrastructure

We ran the encoding models on a NVIDIA RTX A5000 GPU. The average training runtime was around 150 minutes. We used the himalaya library (la Tour et al., 2022). The hyperparameters are searched over 1000 iterations.

## B Layer-wise comparison of predictions from BERT and GPT-2 hidden state embeddings

In order to determine in which layer of BERT the hidden states produced the best predictions of brain responses, we estimated encoding models for each layer. We chose the 8th layer as it performed best (see Figure 6). We ran the same models with the hidden states of GPT-2.

**Figure 6:**
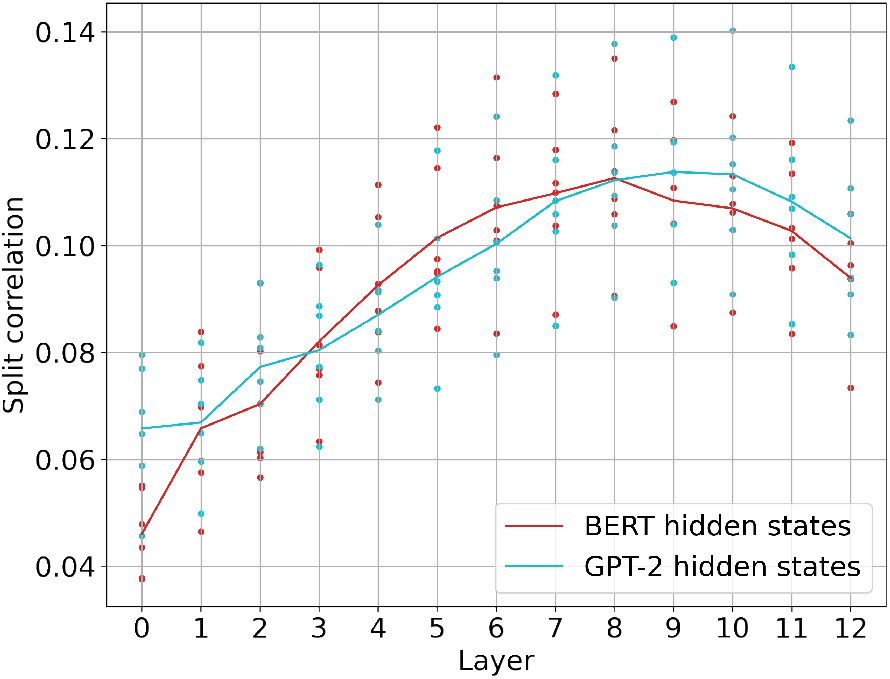
Layer-wise comparison of prediction performance of contextual embeddings extracted from BERT and GPT-2 hidden states. The average performance across participants with BERT embeddings and GPT-2 embeddings are shown with the red and blue lines, respectively. Each dot represents a participant. This comparison led us to choose the hidden states from layer 8 of BERT as contextual embeddings for our experiments.

## C Cortical maps of other participants

Figure 7 through Figure 12 show cortical maps for additional models and other participants. These figures are analogous to Figure 2 and Figure 5.

**Figure 7:**
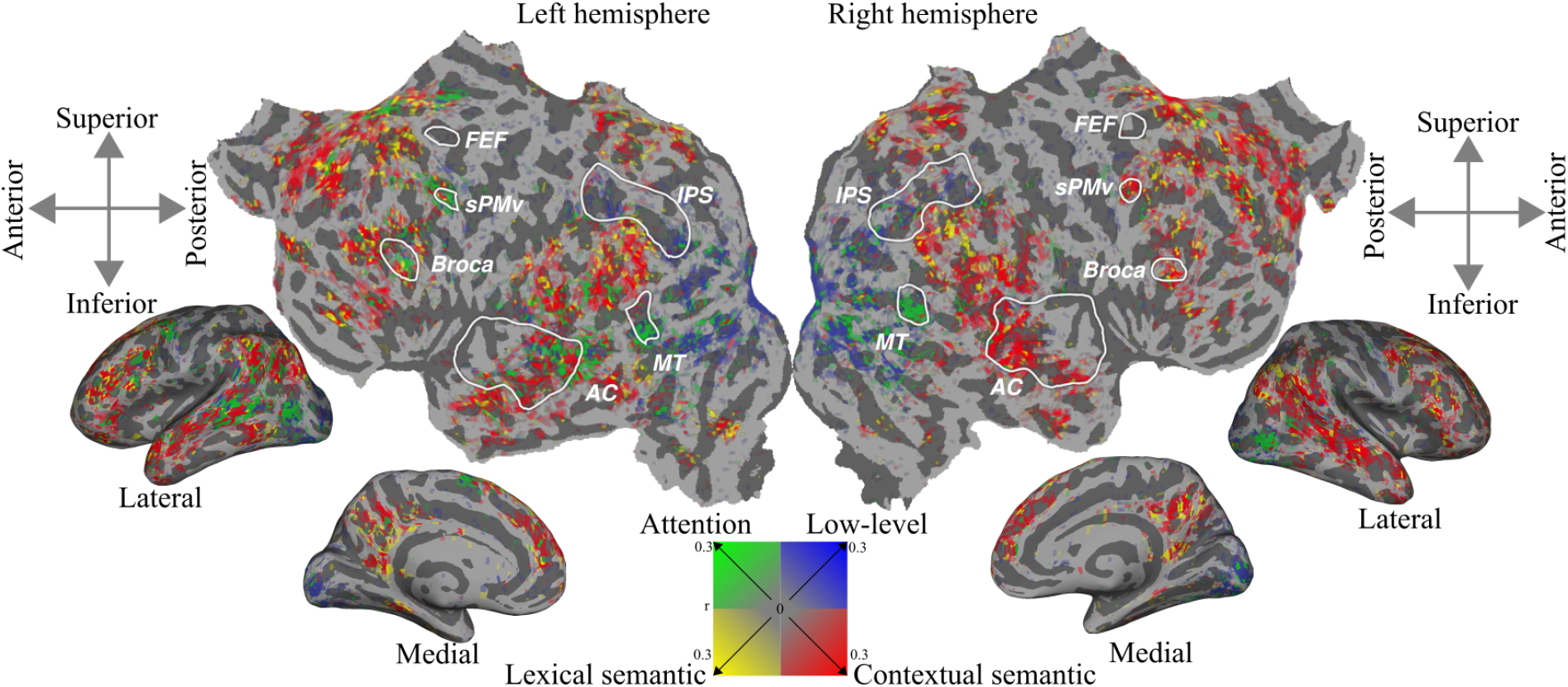
Prediction performance of encoding model with BERT attention weights, low-level features, lexical embeddings, and contextual embeddings across the cerebral cortex for participant 1.

**Figure 8:**
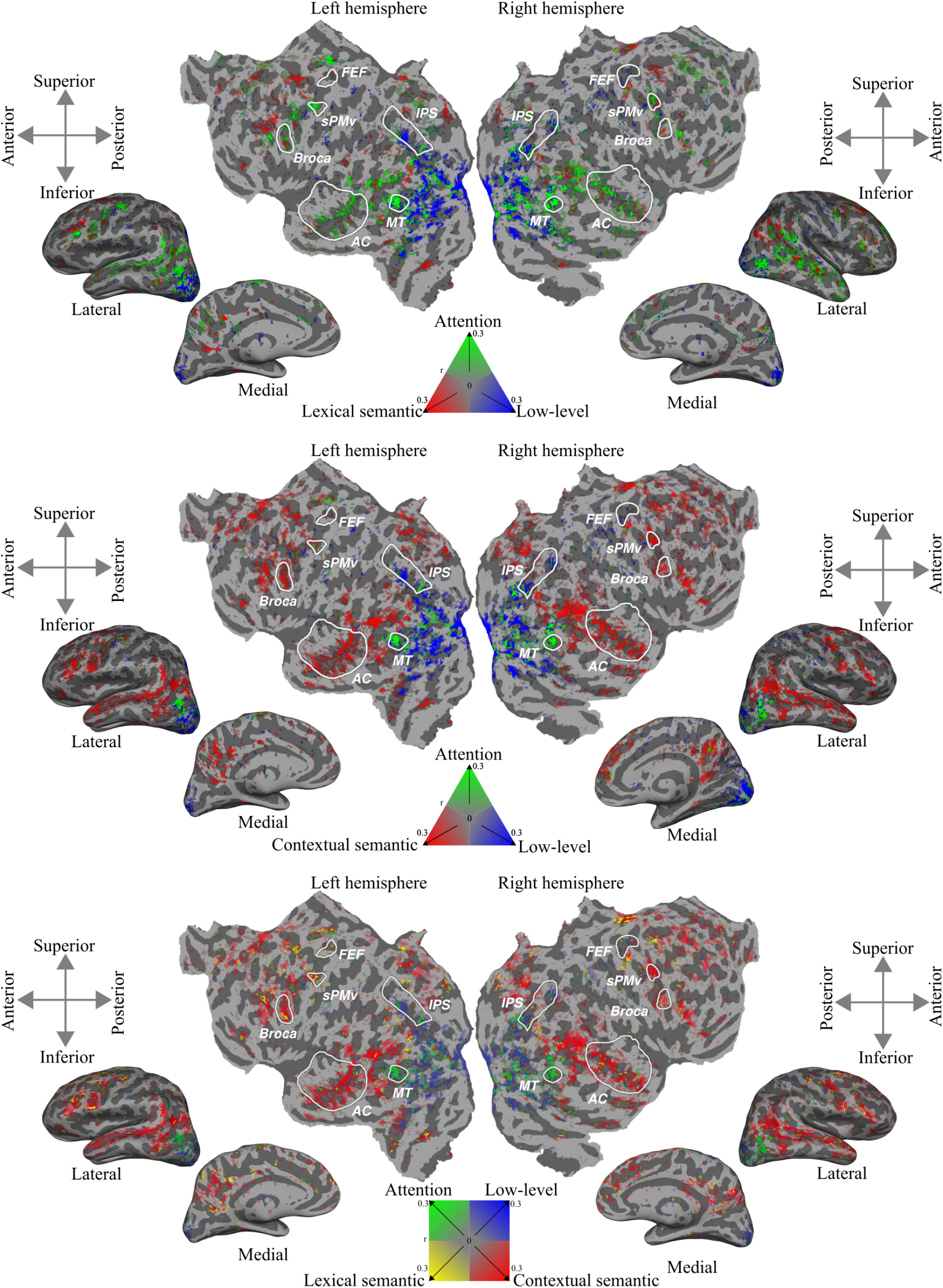
Prediction performance of several encoding models across the cortex for participant 2. (top) Encoding model estimated with BERT attention weights, low-level features and lexical embeddings. (middle) Encoding model estimated with BERT attention weights, low-level features and contextual embeddings. (bottom) Encoding model estimated with BERT attention weights, low-level features, lexical embeddings, and contextual embeddings.

**Figure 9:**
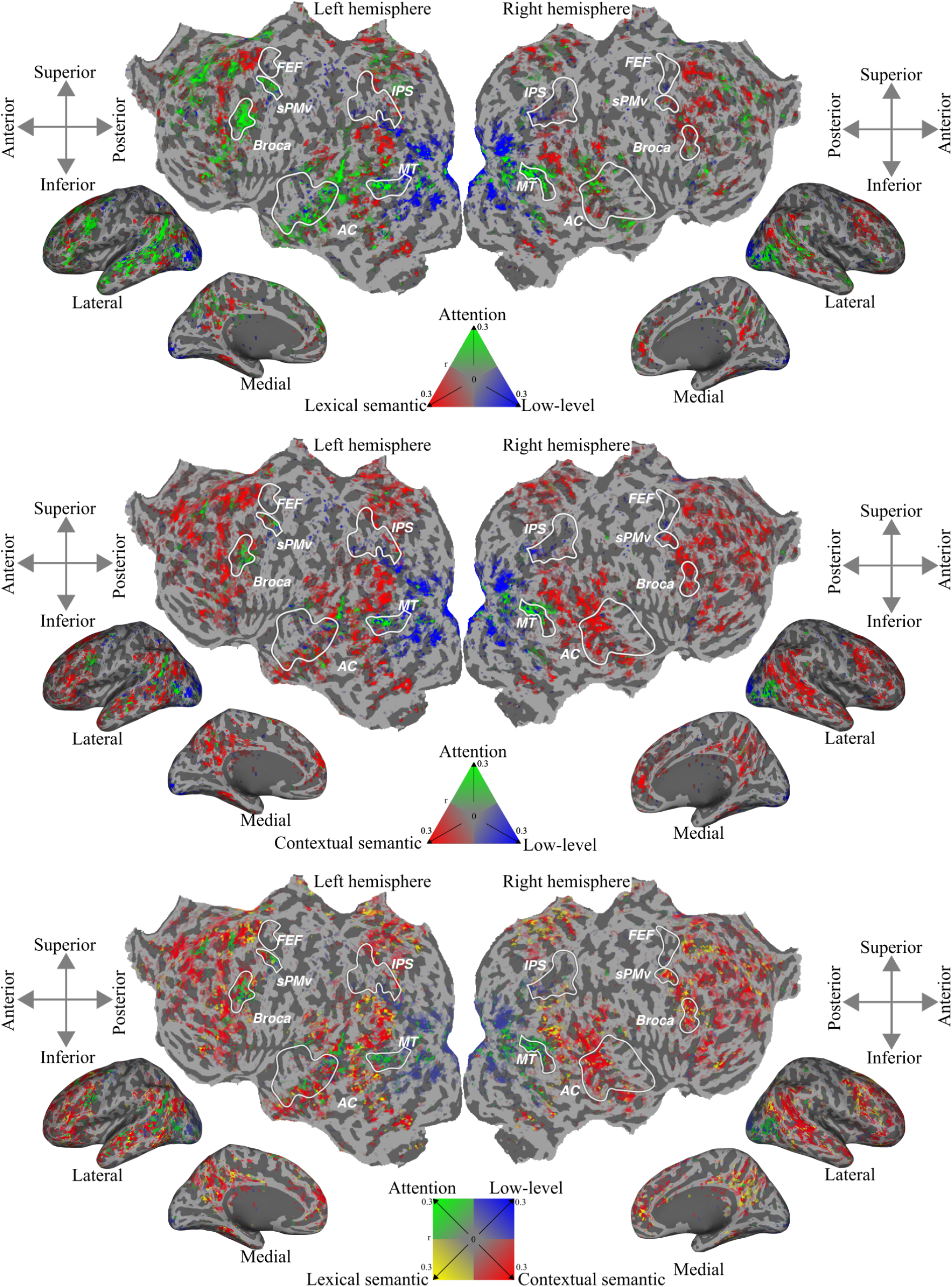
Prediction performance of several encoding models across the cortex for participant 3. (top) Encoding model estimated with BERT attention weights, low-level features and lexical embeddings. (middle) Encoding model estimated with BERT attention weights, low-level features and contextual embeddings. (bottom) Encoding model estimated with BERT attention weights, low-level features, lexical embeddings, and contextual embeddings.

**Figure 10:**
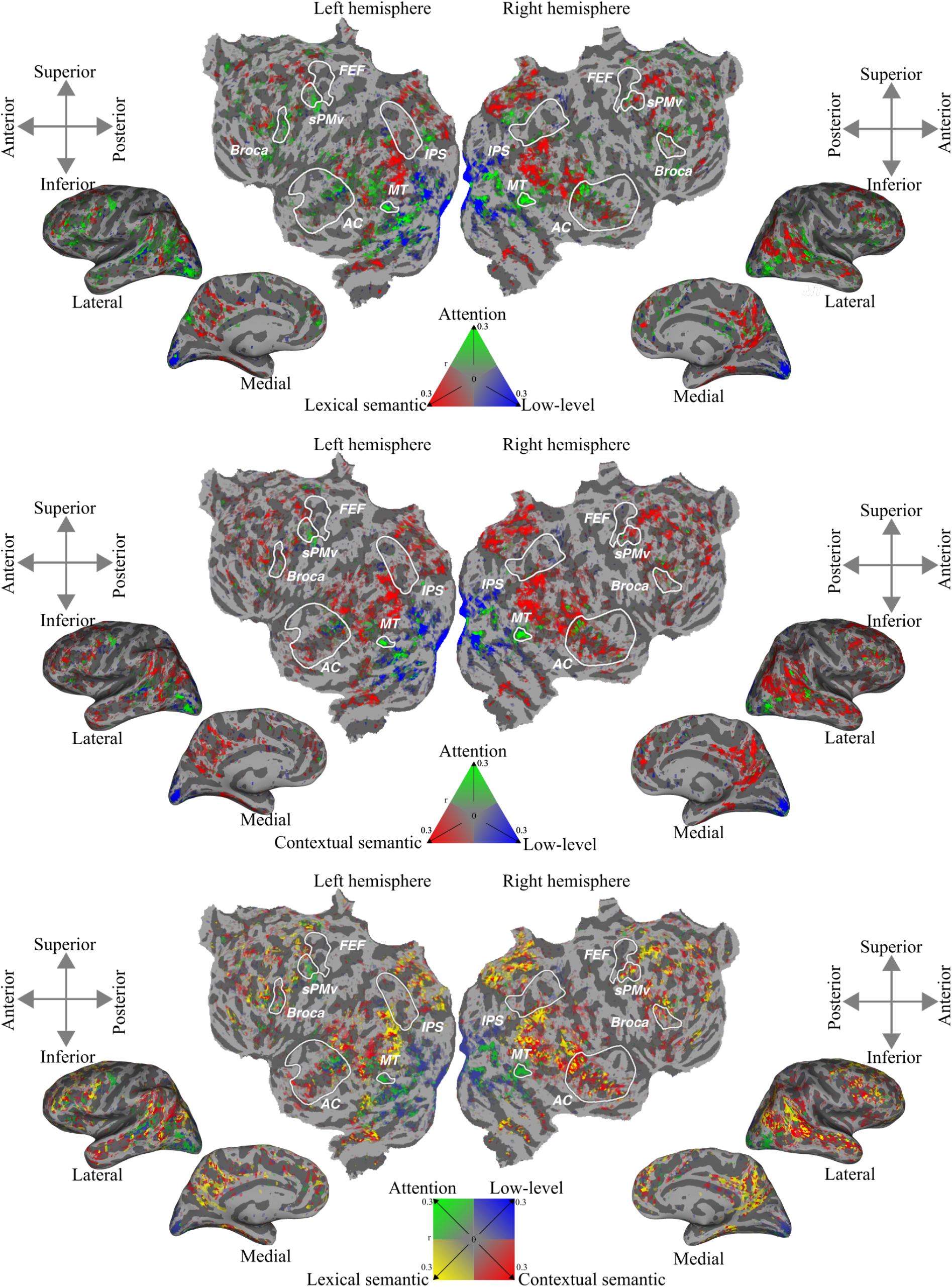
Prediction performance of several encoding models across the cortex for participant 4. (top) Encoding model estimated with BERT attention weights, low-level features and lexical embeddings. (middle) Encoding model estimated with BERT attention weights, low-level features and contextual embeddings. (bottom) Encoding model estimated with BERT attention weights, low-level features, lexical embeddings, and contextual embeddings.

**Figure 11:**
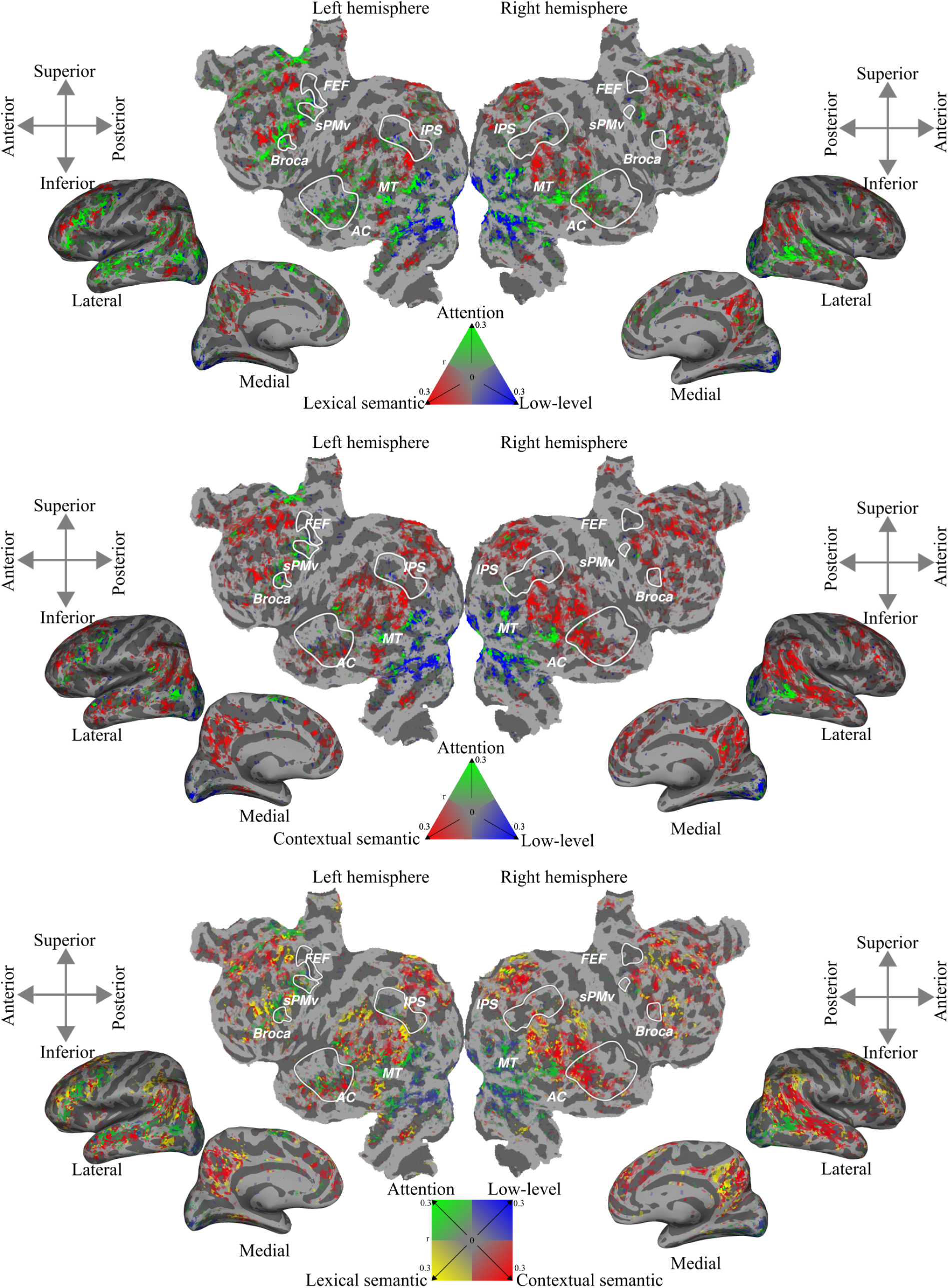
Prediction performance of several encoding models across the cortex for participant 5. (top) Encoding model estimated with BERT attention weights, low-level features and lexical embeddings. (middle) Encoding model estimated with BERT attention weights, low-level features and contextual embeddings. (bottom) Encoding model estimated with BERT attention weights, low-level features, lexical embeddings, and contextual embeddings.

**Figure 12:**
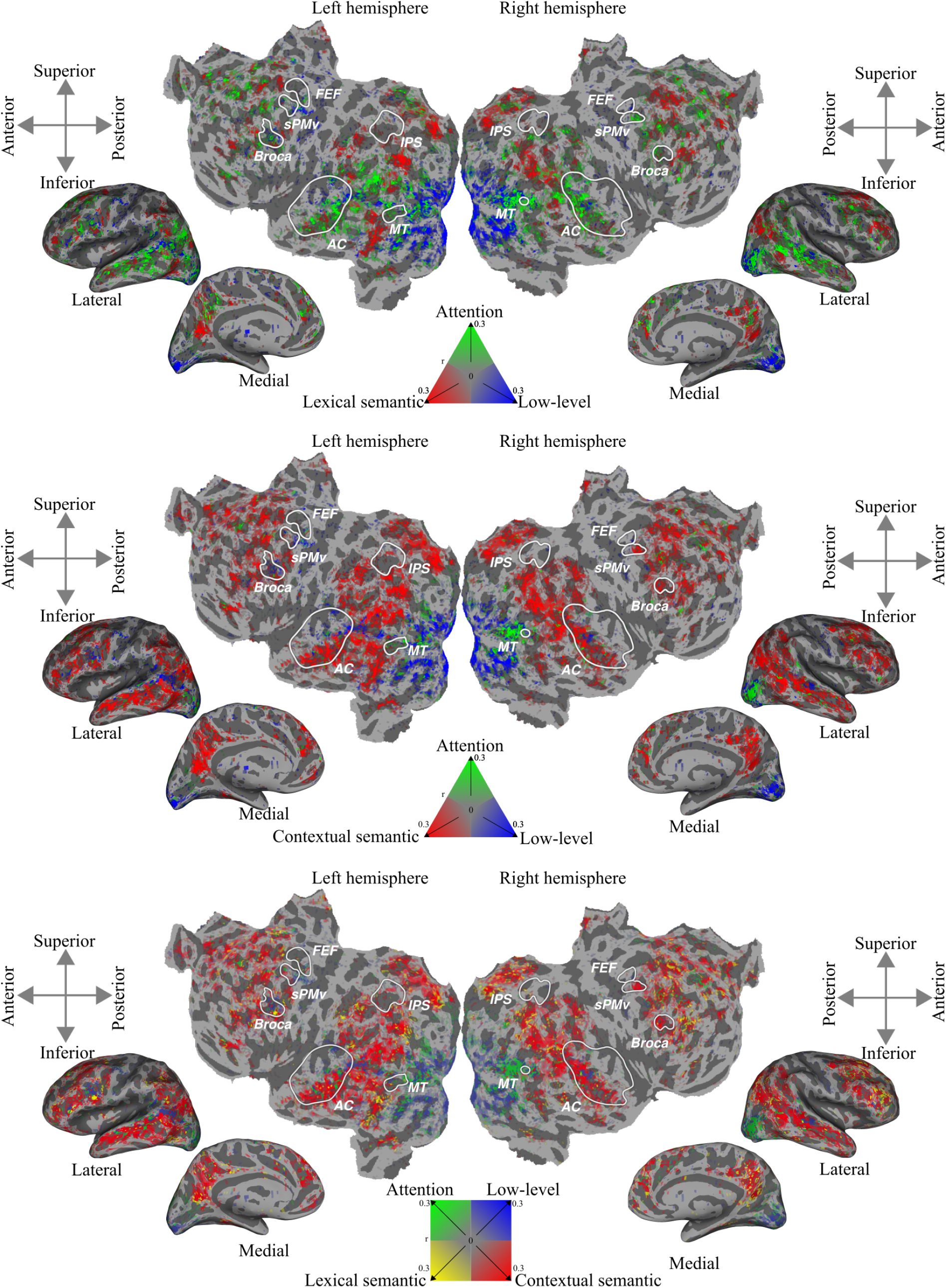
Prediction performance of several encoding models across the cortex for participant 6. (top) Encoding model estimated with BERT attention weights, low-level features and lexical embeddings. (middle) Encoding model estimated with BERT attention weights, low-level features and contextual embeddings. (bottom) Encoding model estimated with BERT attention weights, low-level features, lexical embeddings, and contextual embeddings.

## Footnotes

1 This dataset is openly available: https://berkeley.box.com/v/Deniz-et-al-2019.

## Notes

### Competing Interest Statement

The authors have declared no competing interest.

